## Supplementary figures and images for "A choline-releasing glycerophosphodiesterase essential for phosphatidylcholine biosynthesis and blood stage development in the malaria parasite"

### Supplementary Figure 1

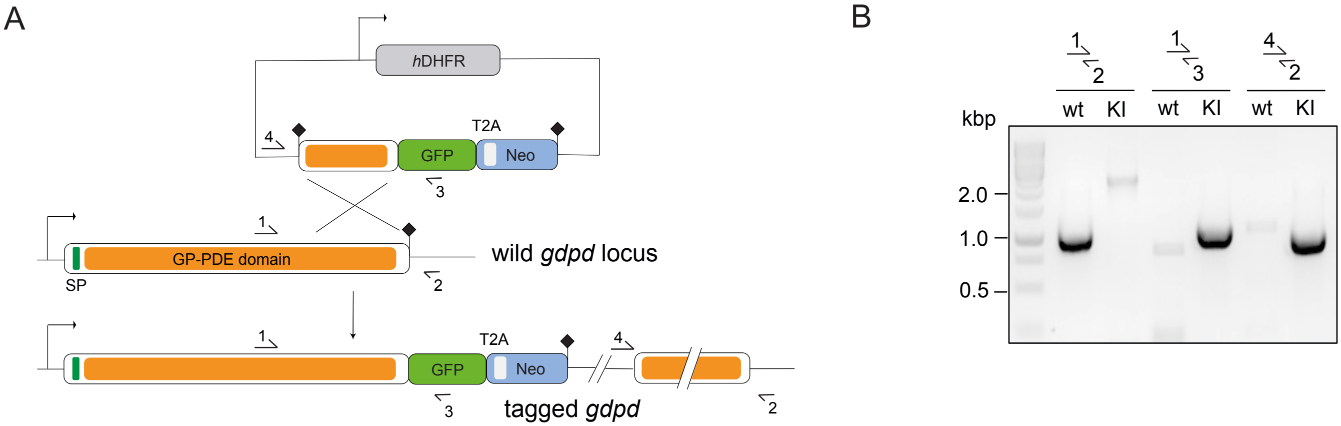

### Supplementary Figure 2

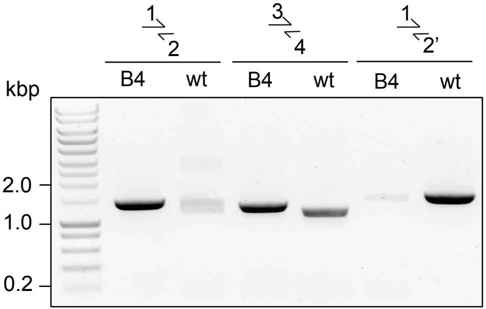

### Supplementary Figure 3

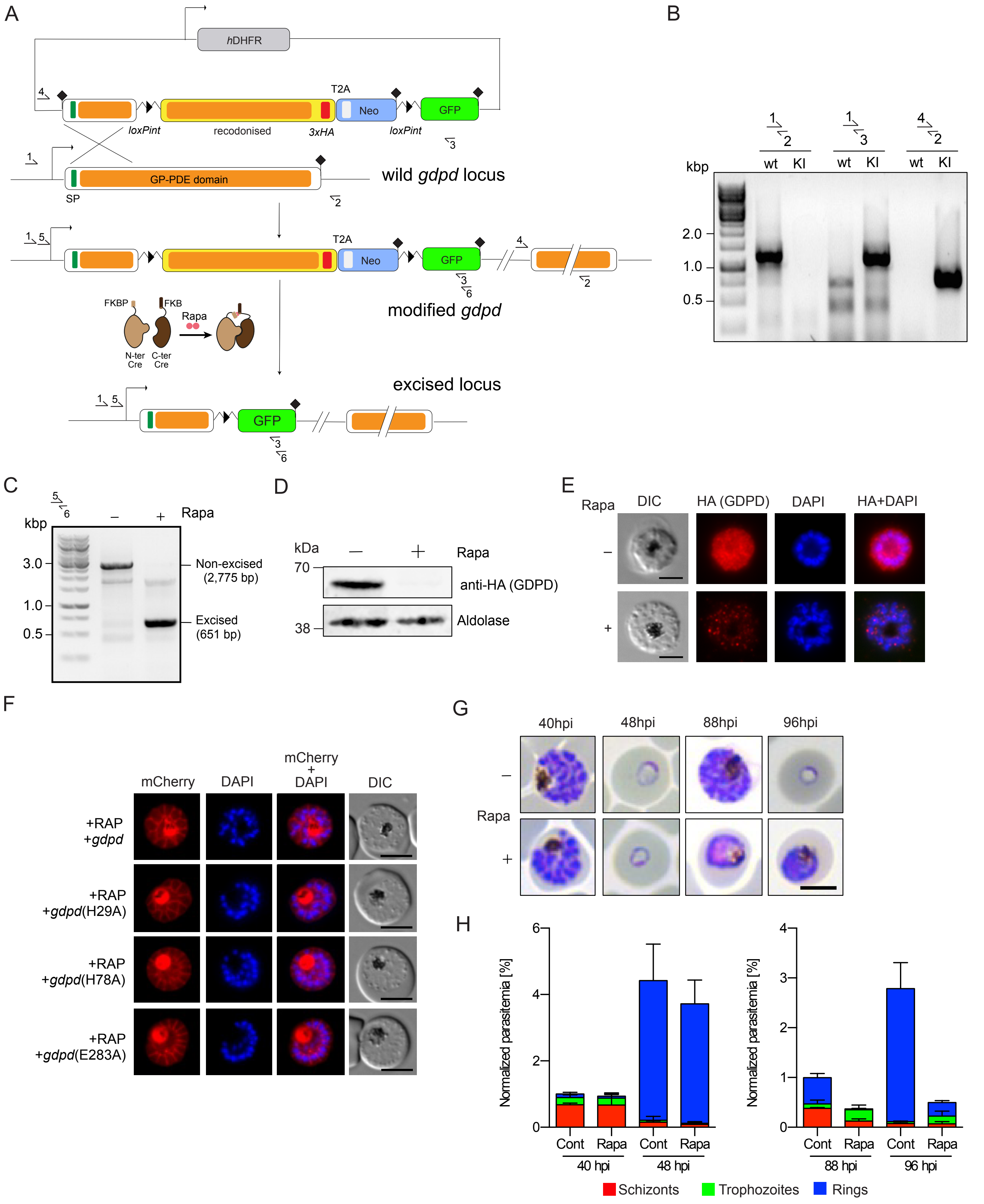

### Supplementary Figure 4

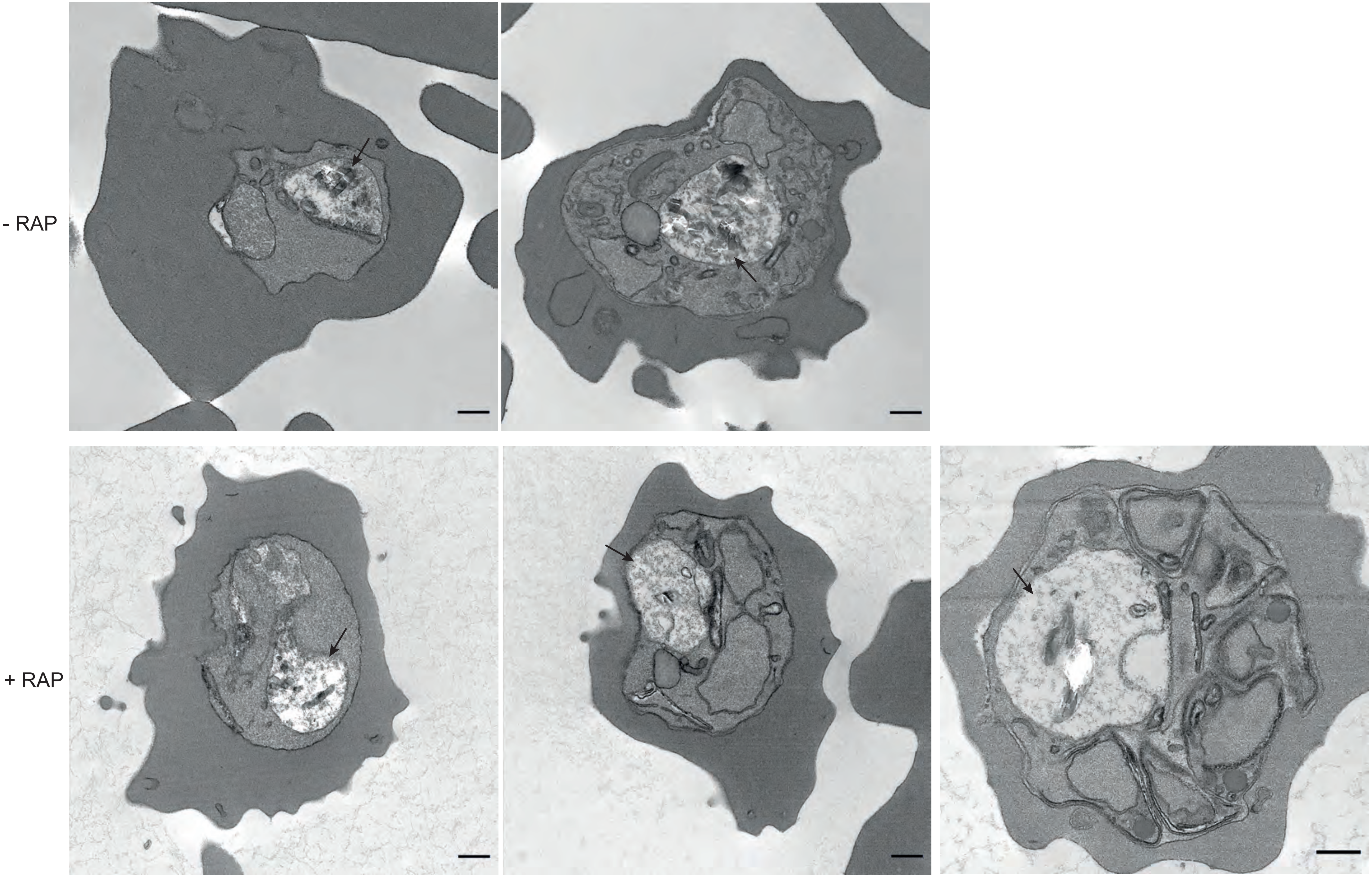

### Supplementary Figure 5

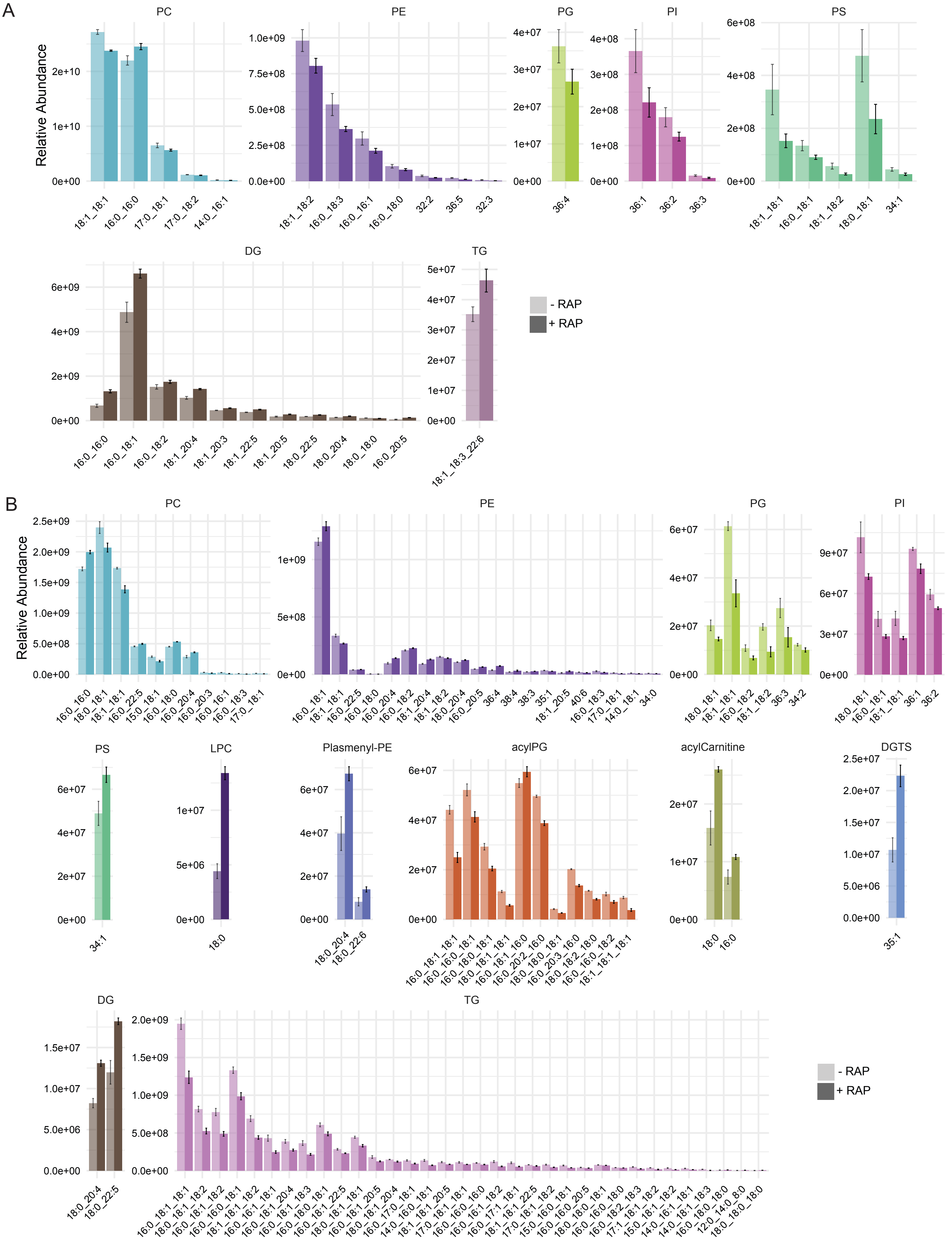

### Supplementary Figure 6

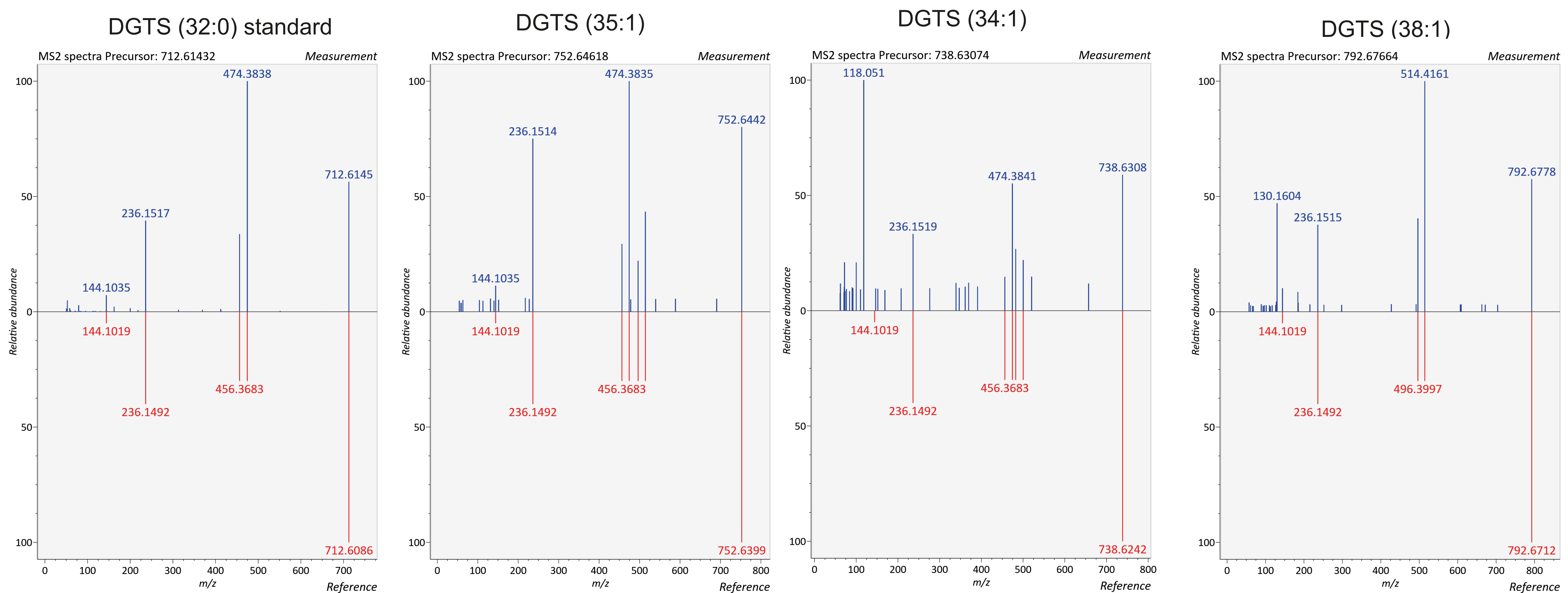

### Supplementary Figure 7

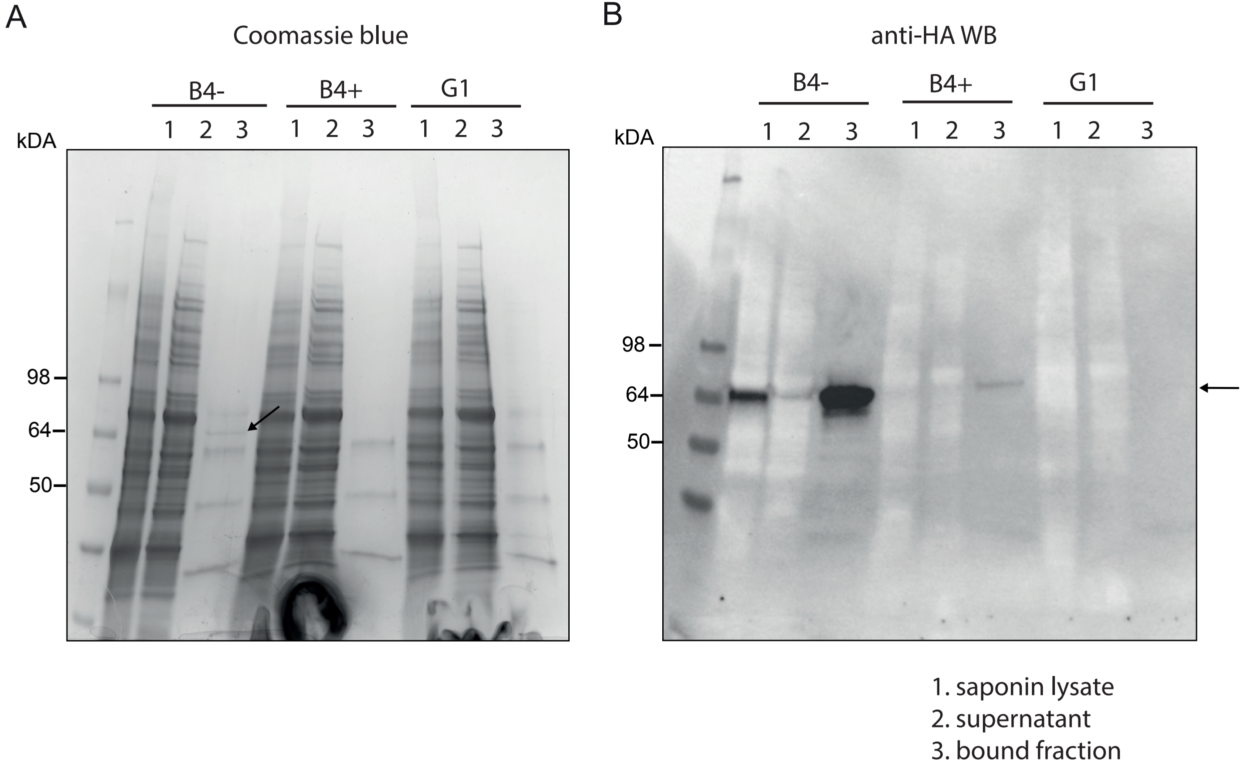

### Supplementary Figure 8

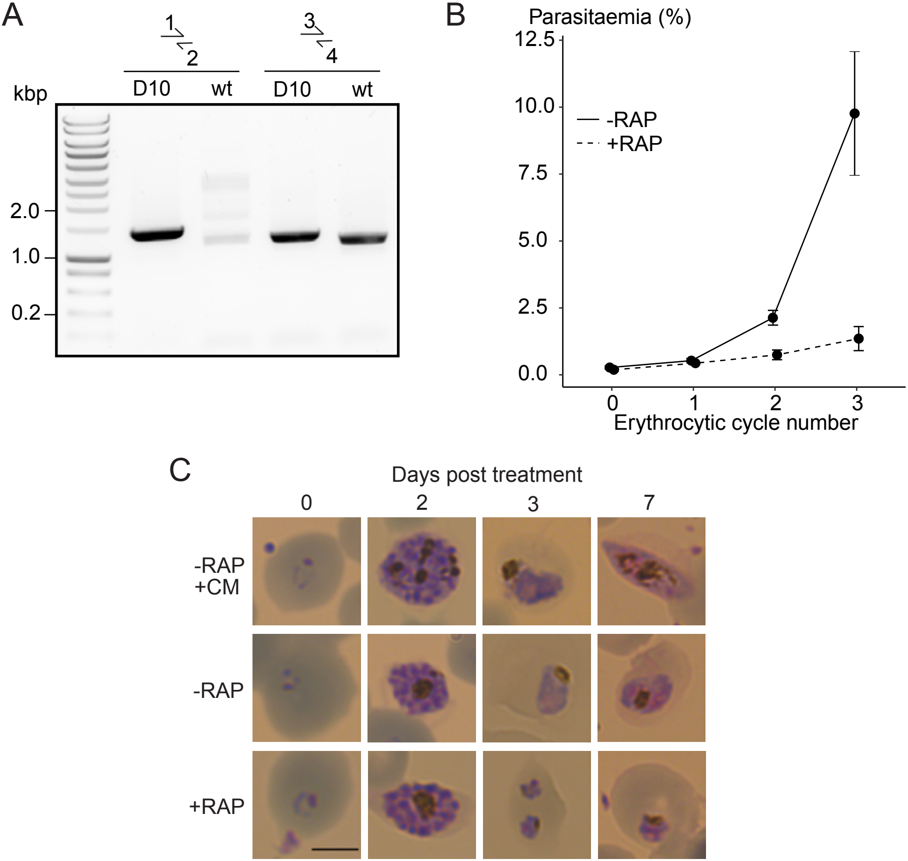

### Supplementary Figure 9

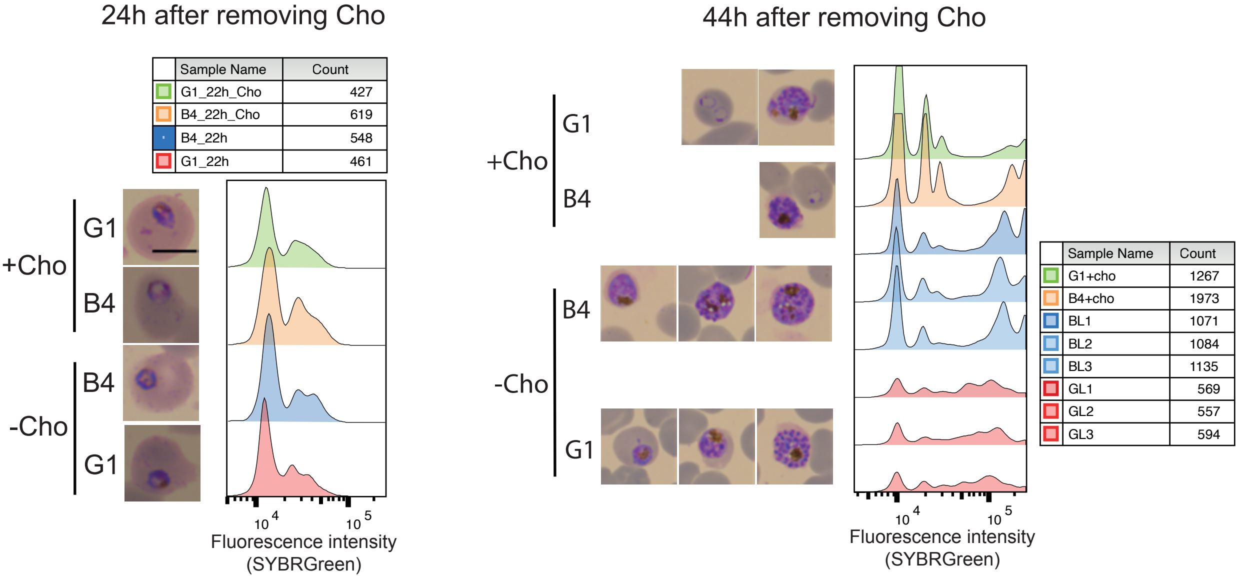

### Supplementary Figure 10

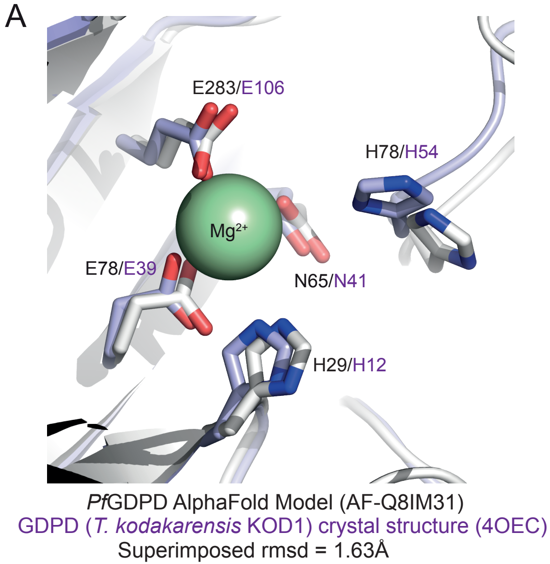


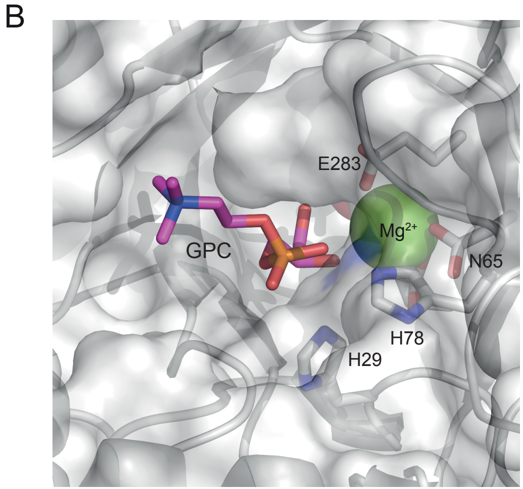


| Ligand | ICM score |
|---|---|
| G3P | -11.38 |
| GPC | -8.16 |
| GPE | -7.94 |
| GPS | -2.84 |
| lysoPC | 4.27 |

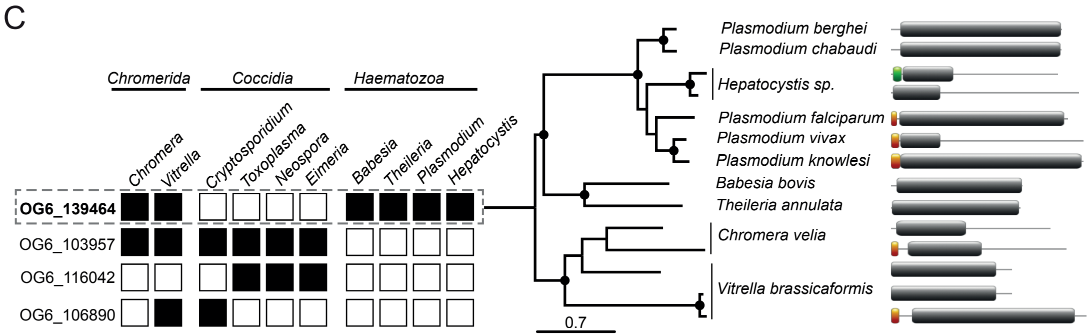
